# Partial elimination of *Wolbachia* in the Asian citrus psyllid, *Diaphorina citri*, increases phytopathogen acquisition and decreases fitness

**DOI:** 10.1101/2023.10.17.562746

**Authors:** Erik L. Roldán, Lukasz Stelinski, Kirsten S. Pelz-Stelinski

## Abstract

*Wolbachia* is a maternally inherited intracellular bacterium that infects a wide range of arthropods. *Wolbachia* can have a significant impact on host biology and development, often due to its effects on reproduction. We investigated *Wolbachia* infection in the Asian citrus psyllid, *Diaphorina citri*, which transmits *Candidatus* Liberibacter asiaticus (*C*Las), the causal agent of citrus greening disease. *D. citri* are naturally infected with *Wolbachia*; therefore, investigating *Wolbachia*-mediated effects on *D. citri* fitness and *C*Las transmission required artificial elimination of this endosymbiont with application of doxycycline. Doxycycline treatment of psyllids reduced *Wolbachia* infection by approximately 60% in both male and female *D. citri*; however, this reduction varied between generations of treated psyllids. Psyllids treated with doxycycline exhibited higher *C*Las acquisition as both adults and nymphs as compared with negative controls. In addition, doxycycline-treated psyllids exhibited decreased fitness as measured by reduced egg and nymph production as well as adult emergence as compared with controls lines where *Wolbachia* was not manipulated. Our results indicate that *Wolbachia* benefits *D. citri* by improving fitness and potentially competes with *C*Las by interfering with phytopathogen acquisition. Targeted manipulation of endosymbionts in this phytopathogen vector may yield disease management tools.

**Importance of the study:** This study provides insights into the critical role of *Wolbachia* in the Asian citrus psyllid, *Diaphorina citri*, a known vector of the presumed causal agent of citrus greening disease. Our data indicate a beneficial role of *Wolbachia* in *D. citri*, whereby the bacteria greatly enhance overall fitness and affect sex ratio of the host but interfere with acquisition of the phytopathogen, *Candidatus* Liberibacter asiaticus (*C*Las). By artificially eliminating *Wolbachia* from psyllids, the study confirms its endosymbiotic role and opens potential avenues for development of disease control methods. Specifically, our data suggest that targeted manipulation of insect endosymbionts like *Wolbachia* may potentially contribute new tools for management of this economically devastating and globally widespread disease of citrus.

## Introduction

*Wolbachia pipientis* is a widespread intracellular bacterium found in a variety of species, including nematodes (Luck et al., 2014; Voronin et al., 2015), insects (Li et al., 2015), and acarids (Glowska et al., 2015). It mainly affects reproduction in hosts (Bourtzis et al., 1996; Werren, 1997), by manipulating host sex via feminization (Asgharian et al., 2014; Kern et al., 2015), male killing (Zeh et al., 2005; Hornett et al., 2014), parthenogenesis (Goryacheva and Andrianov, 2015; He et al., 2015), and cytoplasmic incompatibility (Gerardo and Parker, 2014; Carrington et al., 2015; Ming et al., 2015; Zhang et al., 2015), and can spread rapidly through host populations as a result. Previous research has focused on the mechanism(s) underlying reproductive manipulation of hosts by *Wolbachia* and its potential use to manipulate populations of economically important pests (Curtis and Sinkins 1998, Presgraves 2000, Sinkins and O’Neill 2000, Taylor et al. 2018). *Wolbachia* has also been linked to host speciation and reproductive isolation (Wade, 2001), and is known to decrease fertility, fecundity, and longevity in arthropods (Fleury et al. 2000, Weeks et al. 2001). Some arthropod hosts require *Wolbachia* to produce viable offspring, and their fecundity is reduced when the bacteria are absent (Dedeine et al., 2001). Several studies have generated infected and uninfected isolines of insect hosts to unravel physiological consequences of *Wolbachia* infection (Dobson and Rattanadechakul, 2001).

*Diaphorina citri* Kuwayama (Hemiptera: Liviidae) is a devastating citrus pest because it transmits *Candidatus* Liberibacter asiaticus (*C*Las), a likely causal agent of citrus greening or Huanglongbing (HLB) (Bove 2006, Grafton-Cardwell et al. 2013). HLB is a devastating disease, which causes leaf mottling, asymmetric chlorosis, and yellowing of leaf veins and midribs. Infected trees yield fewer fruit that are smaller, lopsided, bitter, and may retain their green color (Bove 2006). Management of HLB primarily relies on detection of infected plants and insects, calendar-based applications of insecticides and antibiotics, removal of infected plant material, and production of nursery plants under quarantine (Grafton-Cardwell et al., 2013; Stelinski, 2019; Singerman and Rogers, 2020). Current methods have not prevented widespread proliferation of the disease, are unsustainable, and not economically viable; therefore, novel strategies for managing *C*Las transmission by *D. citri* are needed.

*D. citri* harbor three maternally inherited bacterial endosymbionts: *Candidatus* Carsonella ruddii, *Candidatus* Profftella armature, and *Wolbachia pipientis*. Bacterial endosymbionts provide essential nutrients lacking in the diet and aid in food digestion and detoxification. Moreover, endosymbionts play roles in insect defense systems by enhancing pathogen and parasitoid resistance, mediating thermal tolerance of their hosts, and facilitating use of novel hosts (Oliver et al., 2005, 2010; Currie et al., 2006; Dunbar et al., 2007; Hedges et al., 2008; Teixeira et al., 2008; Brownlie and Johnson, 2009; Tsuchida et al., 2011). *Candidatus* Carsonella ruddii (Gammaproteobacteria), a primary obligate endosymbiont, resides in bacteriocytes on the surface of the bacteriome (Thao et al., 2000; Nakabachi et al., 2006, 2013). *Candidatus* Profftella armature (Betaproteobacteria) is a secondary facultative endosymbiont and resides in the syncytial cytoplasm of the bacteriome (Nakabachi et al., 2013). Profftella contributes defense mechanisms to *D. citri* and may facilitate transmission of *C*Las (Ramsey et al., 2015). *Wolbachia* (Alphaproteobacteria) is a well-known arthropod reproductive endosymbiont that infects at least 40% of all arthropod species (Zug and Hammerstein, 2012) and inhabits various *D. citri* tissues including the fat body, salivary gland, ovaries, nervous system, tracheal cells, and midgut (Ammar et al., 2011a, 2011b; Kruse et al., 2017). *Wolbachia* can induce cytoplasmic incompatibility (CI) and enhance host fitness; however, eliminating *Wolbachia* can reduce egg production (Dong et al., 2006). Several studies have observed co-localization of *Wolbachia* with *C*Las in the gut cells of infected adult *D. citri*; however, the exact functions of these endosymbionts and their possible effects on acquisition and transmission of *C*Las by *D. citri* remain unclear (Kruse et al., 2017; Mann et al., 2018).

Given the prevalence of *Wolbachia* in infected insect populations, identifying uninfected individuals in natural hosts can be challenging (Dobson and Rattanadechakul, 2001; Chu et al., 2016). Artificial removal of *Wolbachia* using antibiotics is an alternative approach to generating aposymbiotic strains and gaining a better understanding of the range of functions *Wolbachia* serves in its hosts (Dobson and Rattanadechakul, 2001). Tetracyclines are effective in reducing symbiont infection in insects (Dobson and Rattanadechakul, 2001; Ruan et al., 2006; Raina et al., 2015a; Lee et al., 2016). Doxycycline has also demonstrated some efficacy in reducing *Wolbachia* infection in filarial nematodes and decreases populations of *Wolbachia* up to 98.5% after treatment (Rao et al., 2012).

Manipulation of endosymbionts such as *Wolbachia* may be a viable approach to control insect pests (Weinert et al., 2015). The physiological interactions between *Wolbachia* and *D. citri* remain poorly characterized, largely due to technical challenges in culturing the endosymbiont in vitro and obtaining aposymbiotic *D. citri* (Raina et al., 2015a, 2015b). This study addresses the second challenge by eliminating *Wolbachia* from *D. citri*. We investigated whether *Wolbachia* alters *C*Las transmission by *D. citri* and how *Wolbachia* infection affects the developmental timing, longevity, and reproduction of this phytopathogen vector.

## Material and Methods

### Insect Culture

*D. citri* infected with *Wolbachia* were obtained from a *C*Las-free culture maintained on *Bergera koenigii* (L.) (Rutaceae) in secured greenhouses at the Citrus Research and Education Center (CREC) in Lake Alfred, FL (Chu et al., 2017). Cultures were maintained at 27 °C and 65% relative humidity and a 14:10 h (L:D) photoperiod. Psyllids were confirmed negative for *C*Las as determined by qPCR (described below) every two to three months.

### Antibiotic Selection

Mortality assays were conducted with *D. citri* to optimize treatment dosage for doxycycline (Sigma-Aldrich, St. Louis, Missouri, USA), tetracycline (Sigma-Aldrich, St. Louis, Missouri, USA), and rifampicin (Fisher Bioreagents, Pittsburgh, Pennsylvania, USA). Three concentrations (10 mg/ml, 5 mg/ml, and 2.5 mg/ml) of each antibiotic were evaluated and dissolved in deionized water, except for rifampicin which was dissolved in 5% dimethyl sulfoxide (DMSO) deionized water. Adults (3-5 days old) were injected under the prothoracic sclerite with an antibiotic solution (0.14 μL) and transferred to a *B. koenigii* plant for 20 days. Microinjections were performed using a CO_2_ powered injector at five psi with a borosilicate glass capillary (World Precision Instruments, Sarasota, FL, USA). The borosilicate glass capillary was custom designed to 0.8mm using a PUL-1000 (World Precision Instruments, Sarasota, FL, USA).

A subsequent experiment was conducted to determine the most effective doxycycline concentration and treatment duration for eliminating *Wolbachia* infection. Adults (15 female, 15 male) were injected with three doxycycline concentrations (10 mg/ml, 5 mg/ml, 2.5 mg/ml) and then transferred to separate *B. koenigii* plants to prevent mating. *Wolbachia* infection was assessed by subsampling three adults from each concentration after 24, 48, and 72 hours. The experiment was repeated five times.

### Doxycycline Treatments

Newly emerged (0–1-day-old) *D. citri* adults (60 male : female pairs) were microinjected with 10 mg/ml of doxycycline solution. This cohort is referred to as ‘F0’. Then, adults were separated by sex and placed onto different *B. koenigii* plants to prevent mating. After 24 hours, 15 separate pairs of male and female *D. citri* adults were collected and confined as pairs within fine nylon mesh sleeves enveloping newly unfurled leaf flush to allow mating on four individual plants. Forty of the initial F0 psyllids were collected at 30 and 45 d after microinjection for detection of *Wolbachia* by qPCR, as described below. The F0 adults were discarded, and F1 offspring obtained from the injected mating pairs were reared on *B. koenigii* until the adult stage, then injected with 10 mg/ml of doxycycline, and returned to plants. The above procedure was repeated to obtain several continuous generations of doxycycline-injected psyllids (F2-F9). A control group of psyllids was similarly generated; however, it was injected with deionized water instead of the antibiotic solution. The numbers of females and males were quantified per generation to determine sex ratios.

### Mortality, Longevity, and Reproduction

The objective of this experiment was to determine the effect of antibiotic treatment on adult psyllid longevity. Twenty male : female pairs of newly-emerged (0–1-day-old) adults were injected with doxycycline or deionized water (control), as described above, and released onto *B. koenigii*. Pairs of treated psyllids were confined on leaves within mesh cages on individual plants maintained in a climate-controlled environmental chamber at 27 ± 1°C, 65 ± 5% relative humidity, with a 14:10 h light : dark photoperiod. Psyllid mortality was recorded daily for 20 days.

Egg laying was quantified during the first seven days after treatment injection. Female *D. citri* only lay eggs on newly unfurled (0-3 day after budbreak) leaf flush (Tsai and Liu, 2000; Nava et al., 2007). Therefore, the 20 pairs of male : female adults were transferred to newly unfurled flush throughout the fecundity observations to facilitate continuous egg laying. The total number of eggs laid per leaf flush shoot was counted daily under a stereomicroscope for the initial seven days. The number of fertile eggs was determined by quantifying psyllid nymph emergence daily at 7 to 14 days after treatment injections. The same procedure was repeated during generations F0-F9.

### Acquisition Assays

The objective of these experiments was to determine whether reducing populations of *Wolbachia pipientis* within *D. citri* by antibiotic treatment affects acquisition of the *C*Las bacterium. For acquisition experiments, *D. citri* were injected with doxycycline or the control as described above and immediately placed onto *C*Las-infected citrus trees as described by Pelz-Stelinski et al. (2010). Pairs of *D. citri* adult males and females (n = 10) from each treatment group were enclosed within fine nylon mesh sleeves on newly unfurled leaf flush of 2 yr old and *C*Las-infected ‘Valencia’ sweet orange, *Citrus sinensis* (L.) Osbeck (1.20–1.50 m in height), plants. Psyllids caged on plants were allowed a seven-day acquisition access period (AAP). Citrus plants used in this assay were confirmed *C*Las positive by qPCR (CT value between 19-22); however, they were not yet declining due to disease expression at 8-14 months post graft inoculation with pathogen. Thereafter, adults were collected and preserved in 80% ethanol and placed at −20°C until DNA extraction for detection of *C*Las and *Wolbachia*. Eggs were left on trees within mesh sleeves to evaluate *C*Las acquisition by F1 nymphs after emergence. After the *D. citri* nymphs reached the adult stage (ca. two weeks), they were collected to quantify *C*Las acquisition. The assay was replicated ten times on separate *C*Las-infected trees for both doxycycline-treated and control lines. No *C*Las acquisition was observed from uninfected trees run as a negative control (data not shown).

### *D. citri* Endosymbiont Detection

Total DNA was extracted from *D. citri* using the Qiagen DNeasy Blood and Tissue Kit (Qiagen Inc., Valencia, California, USA) according to the manufacturer’s protocol with modifications for isolation of bacterial DNA from arthropods (Pelz-Stelinski et al., 2010). DNA concentration was determined using a NanoDrop 2000 spectrophotometer (Thermo Fisher Scientific, Lafayette, Colorado, USA) and 15 ng was used per PCR reaction. Quantitative real-time polymerase chain reaction (qPCR) analysis was performed using the Applied Biosystems QuantStudio™ 6 Flex Real-Time PCR System (Thermo Fisher Scientific Inc., Waltham, MA, USA). Analysis of endosymbiont DNA was performed using the SYBR Green PCR Master Mix (Life Technologies, LTD, Woolston Warrington, UK) and primer sequences as described in Chu et al. (2016) in a final reaction volume of 19 μL (Table 1). All qPCR reactions were performed in duplicate and qPCR settings consisted of an initial denaturation step of 95°C for 10 min, 40 cycles of 95°C for 15 seconds and 58°C (*Wolbachia*, Carsonella, and Profftella) or 60°C (*wingless*) for 30 s, and a final cycle of 72°C for 30 s. The relative abundance (ΔΔ-CT) of endosymbionts in *D. citri* was determined based on the CT-values of *Wolbachia pipientis, Candidatus* Carsonella ruddii, *Candidatus* Profftella armature, and *wingless* genes, (Livak and Schmittgen, 2001). Using CT-values for *C*Las *16S rRNA*, *wingless*, and *COX* genes, the relative abundance of *C*Las in *D. citri* and plants was determined for the acquisition assays.

**Table 1.**
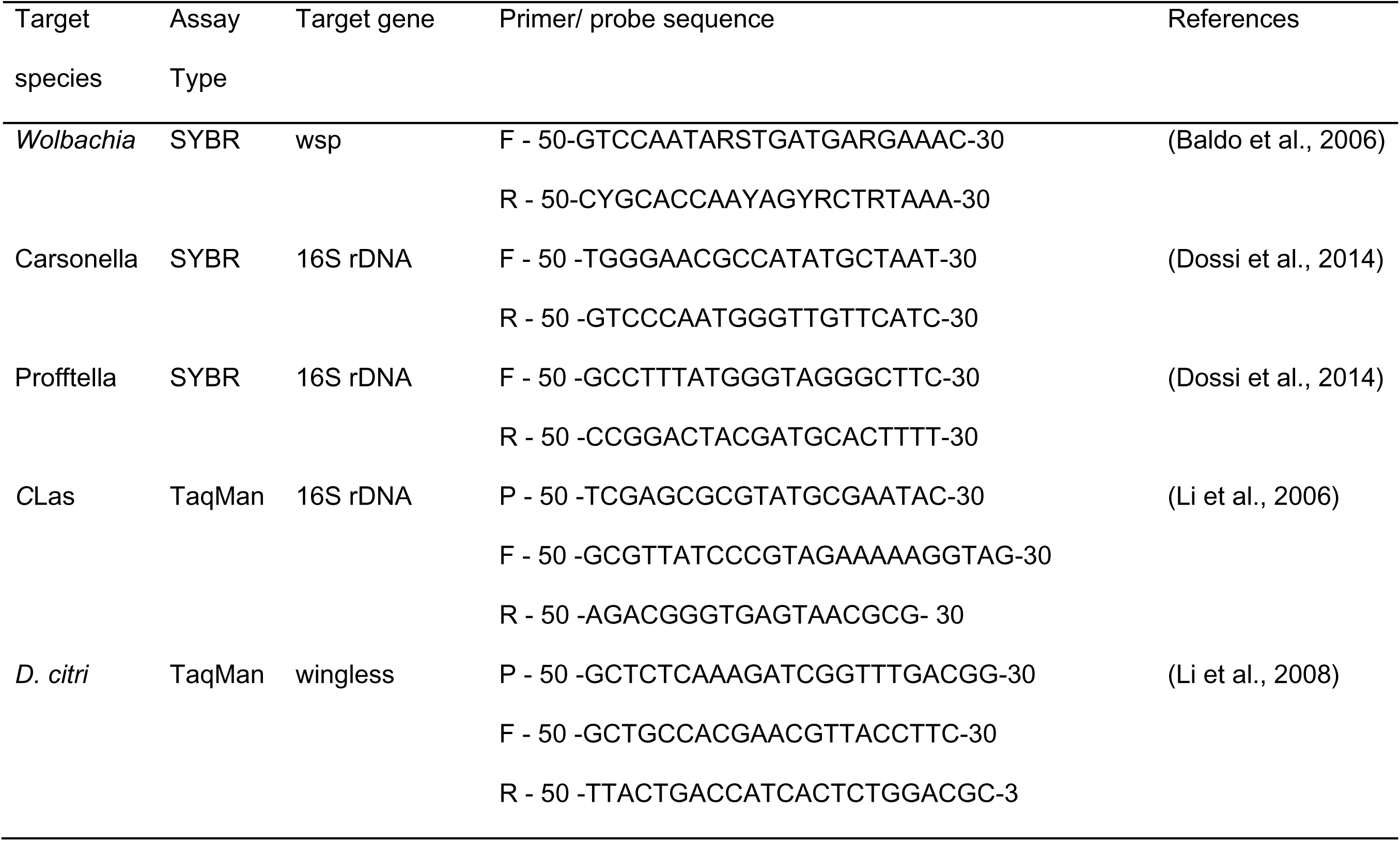
List of primers and probes used to detect *Diaphorina citri* endosymbionts with quantitative PCR.

### Detection of *C*Las in *D. citri* and Citrus Plants

DNA from insect samples and plants was isolated using the DNeasy Blood and Tissue or DNeasy plant kits (Qiagen, Valencia, CA, USA), according to the manufacturer’s instructions. Samples were quantified using a NanoDrop 2000 spectrophotometer (Thermo Fisher Scientific Inc., Waltham, MA, USA). DNA was diluted to *C*Las-specific 16S rDNA from 10 ng μL^−1^ for plant template DNA and 15 ng μL^−1^ for *D. citri* DNA for subsequent quantitative real-time PCR analysis.

Detection of *C*Las in *D. citri* and citrus plants was conducted using multiplex TaqMan real-time PCR methods (Table 1) (Li et al., 2006, 2008). Endogenous control genes *wingless* (*D. citri*) and *cytochrome oxidase* (plants) were selected for the assays, as per Li et al. (Li et al., 2008). The qPCR assays were conducted in triplicate with an ABI 7500 Real-Time PCR System (Applied Biosystems, Waltham, MA, USA). For plant DNA, each reaction mixture was composed of 12.5 μL of PerfeCTaq PCR ToughMix, Low ROX (Quanta BioSciences Inc., Gaithersburg, Maryland, USA), 3 and 3.75 μL of *C*Las and *COX* primers respectively, 3.75 μL of each probe, and 1 μL of DNA template (10ng) in a final volume of 27.75 μL. For *D. citri* DNA, each reaction mixture was composed of 12.5 μL of PerfeCTaq PCR ToughMix, 3 μL of each primer, 3 μL of each probe, and 1 μL of DNA template (15ng) in a final volume of 25.5 μL. Reactions were considered positive for either target sequence if the cycle quantification (CT) value, determined by the ABI 7500 Real-Time software (version 1.4, Applied Biosystems), was ≤ 36. Final primer concentrations were 0.2 μM for *C*Las, *wingless*, and *COX*. Final probe concentrations were 0.1 μM for *C*Las, wingless, and *COX*. Real-time PCR reactions consisted of an initial denaturation step of 95°C for 10 min followed by 40 cycles of 95°C for 15 s for *cytochrome oxidase* and 60°C for 60 s for *wingless*.

## Data Analysis

Mean abundance of endosymbiont DNA between doxycycline- and control-treated psyllids were compared with T-tests. T-tests were also used to compare the mean numbers of eggs laid, as well as the number of psyllid nymphs and adults emerging between the two treatment lines. Survival rates were assessed using the Kaplan-Meier analysis and statistically compared using the log-rank tests. Egg hatch, adult emergence, and sex ratios among antibiotic-injected and control treatments were compared using chi-square analyses. Relative abundance of *C*Las acquired by *D. citri* caged on *C*Las-infected citrus trees was subjected to analysis of variance (ANOVA) followed by post-hoc multiple comparisons (Tukey’s HSD test) for mean separation. The mean *C*Las abundance in *D. citri* and plants was log10(x+0.1) transformed to meet the assumptions of normality. *C*Las infection rates between doxycycline- and control-treated *D. citri* adults and emerging nymphs were compared using chi-square analyses. Relationships between mean *Wolbachia* infection per generation, insect reproductive fitness variables, and endosymbiont titers were investigated using Pearson correlations. Furthermore, correlations between *D. citri* endosymbionts and *C*Las infection were evaluated using insects from acquisition assays. In all cases, the significance level was set at α < 0.05. All statistical analyses were performed using R Studio (R Core Team, 2013).

## Results

### Antibiotic Selection

Antibiotics caused differential effects on *D. citri* survival and endosymbiont population levels. Survival decreased significantly among insects fed rifampicin with higher mortality occurring approximately three days after feeding (Log-rank test, χ^2^(1) = 21.45, P < 0.001). In contrast, *D. citri* survived more than five days after feeding on tetracycline (Log-rank test, χ^2^(1) = 0.78, P = 0.38) or doxycycline (Log-rank test, χ^2^(1) = 0.70, P = 0.40), even using the highest dosage. Survival of *D. citri* declined significantly five days after injection with tetracycline, but with not doxycycline, compared with the control group (Fig. 1). *D. citri* injected with doxycycline survived for more than ten days (Fig. 1).

**Figure 1.**
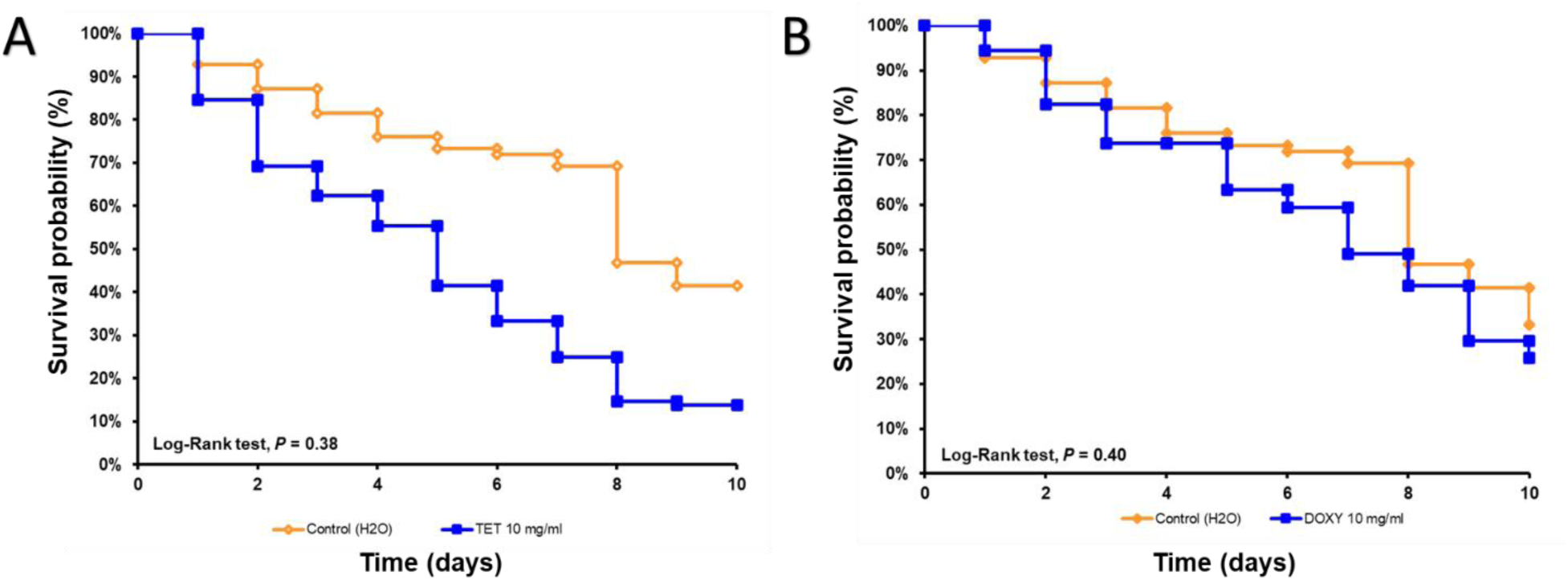
Effect of (A) tetracycline (TET) and (B) doxycycline (DOX) treatment on survival of *Diaphorina citri* as displayed by Kaplan-Meier longevity curves. (A-B) Log-rank test, P < 0.05.

### Susceptibility of *D. citri* Endosymbionts to Doxycycline

To determine the susceptibility of endosymbionts to doxycycline, the abundance of symbionts within psyllids was compared following antibiotic treatment over nine generations of doxycycline-treated and control lines. Abundance of *Wolbachia* was significantly lower in doxycycline-treated males (T-Test, T = 6.66, df = 18, P < 0.01; 54% reduction) and females (T-Test, T = 7.75, df = 18, P < 0.01; 67% reduction) than in respective control lines (Fig. 2A).

**Figure 2.**
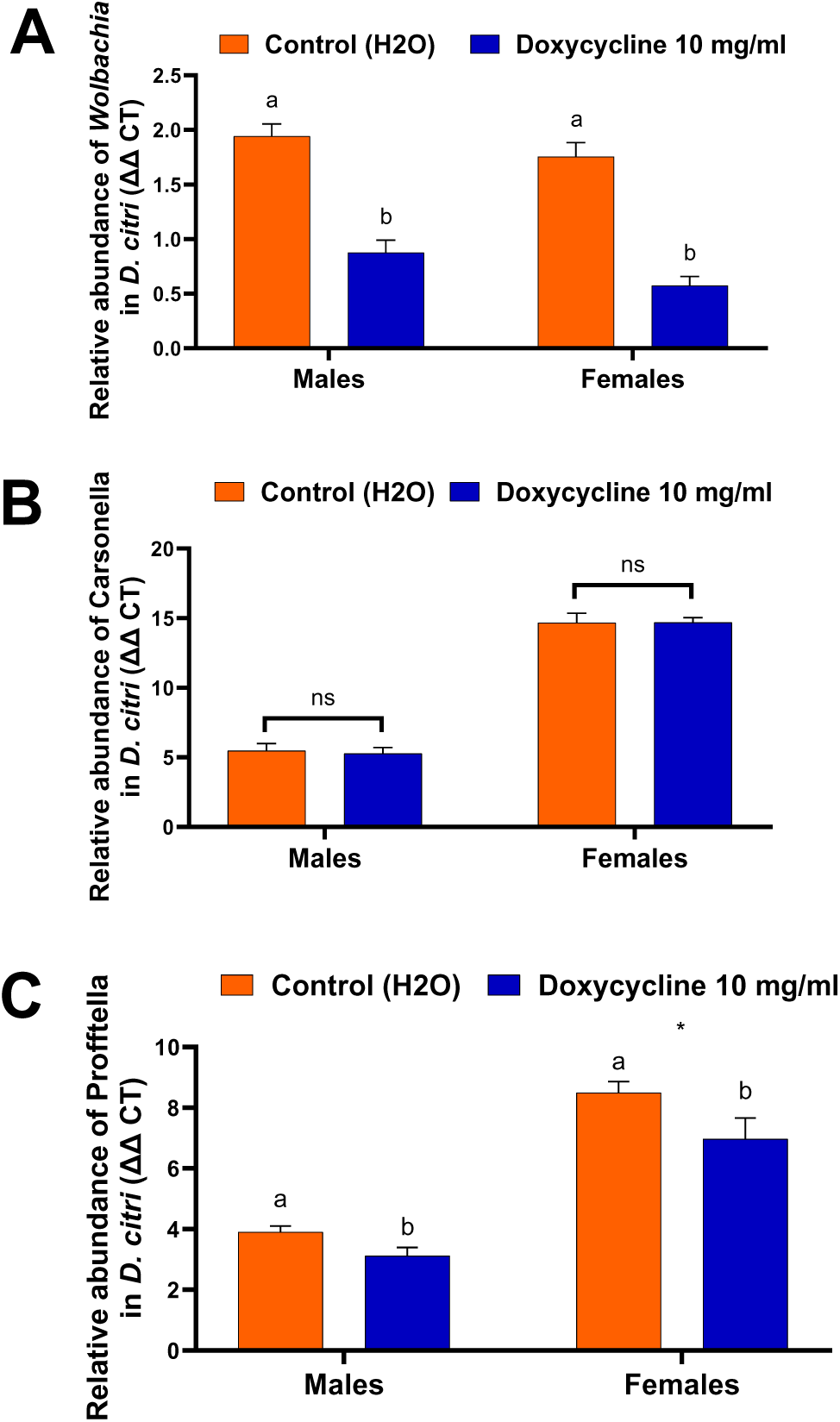
Effect of doxycycline treatment on (A) *Wolbachia,* (B) Proftella, and (C) Carsonella abundance in *Diaphorina citri* males and females. Treatments indicated by “ns” are not significantly different (A-C) T-Test; P < 0.05; * = P < 0.1, Tukey’s HSD).

Doxycycline treatment did not cause a statistical reduction in populations of Carsonella in *D. citri* males (T-Test, T = 0.29, df = 18, P = 0.77) or females (T-Test, T = 0.03, df = 18, P = 0.97) as compared with the control treatment (Fig. 2B). However, Carsonella was approximately 3-fold more abundant in females than males (Fig. 2B). Also, Carsonella was more abundant overall in both sexes of *D. citri* than the other two endosymbionts quantified (Fig. 2B). Interestingly, abundance of *Wolbachia* in female *D. citri* was positively correlated with abundance of Carsonella (Table 2).

**Table 2.** Correlations among *Diaphorona citri* endosymbiont population abundance in doxycycline-treated and control cultures of *D. citri* across nine consecutive generations.

|  |  |  |  |  |  |  |
| --- | --- | --- | --- | --- | --- | --- |
| Doxycycline-treated line | Mean *Wolbachia* |  |  |  |  |  |
| (All generations) | infection ($\Delta\Delta$ CT) | | | | | |
| **Females *D. citri*** | *correlation (r)* | *R2* | *T* | *df* | *P > F* |  |
| Mean Profftella infection ($\Delta\Delta$ CT) | 0.67 | 0.44 | 2.68 | 9 | 0.025 | |
| Control line |  |  |  |  |  |  |
| (All generations) |  |  |  |  |  |  |
| **Females *D. citri*** | *correlation (r)* | *R2* | *T* | *df* | *P > F* |  |
| Mean Carsonella infection ($\Delta\Delta$ CT) | 0.65 | 0.42 | 2.59 | 9 | 0.029 | |
| Mean Profftella infection ($\Delta\Delta$ CT) | 0.66 | 0.44 | 2.66 | 9 | 0.026 | |

Profftella abundance was significantly lower in the doxycycline-treated *D. citri* males (T-Test, T = 2.21, df = 18, P = 0.03; 20% reduction) and females (T-Test, T = 1.93, df = 18, P = 0.07; 17% reduction) as compared with the control line (Fig. 2C). Furthermore, the abundance of *Wolbachia* in *D. citri* females in both doxycycline-treated and control line lines was positively correlated with abundance of Profftella (Table 2).

### Effect of Doxycycline on Mortality, Longevity, and Reproduction of *D. citri* Fecundity, fertility, and development

There was a significant effect of treatment on the total number of eggs produced by generation of *D. citri*. Overall, the doxycycline-treated *D. citri* line produced significantly (T-Test, T = 9.44, df = 78, P < 0.01) fewer eggs than the control line (39% reduction) (Fig. 3A). Fertility, assessed as the percentage of eggs hatched, was significantly higher in the doxycycline-treated *D. citri* line than in the control line (9.71% higher) (Chi-square, X^2^ = 73.63, df = 1, P < 0.01) (Fig. 3B).

**Figure 3.**
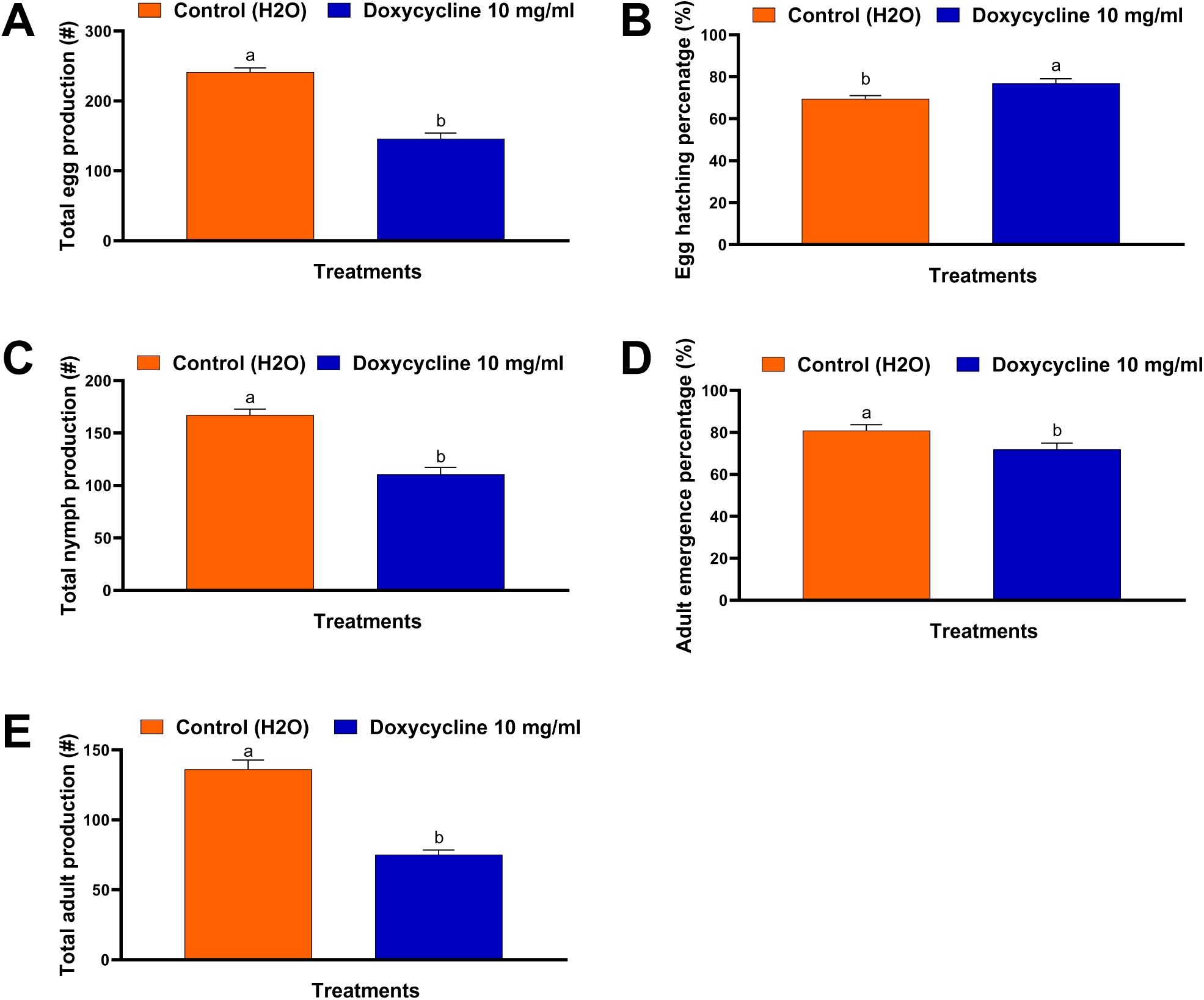
Effect of doxycycline treatment on *Diaphorina citri* (A) total egg production, (B) egg hatch percentage, (C) total nymph production, (D) adult emergence percentage, and (E) the total number of adults produced per generation. Treatments indicated by “ns” are not significantly different. (A, C, D) T-Test P < 0.05; * < 0.1, Tukey HSD; (B, D) Chi-square, P < 0.05.

Doxycycline-treated *D. citri* produced significantly (T-Test, T = 6.42, df = 78, P < 0.01) fewer nymphs than those in the control line (33% decrease) (Figure 3C). Furthermore, emergence of adult *D. citri* was significantly (Chi-square, X^2^ = 246.16, df = 1, P < 0.01) reduced in the doxycycline-treated line than in the control line (8.86% decrease) (Fig. 3D). Furthermore, the control line produced significantly (T-Test, T = 8.14, df = 78, P < 0.01) more adults in total than the doxycycline-treated line (Fig. 2E). *Wolbachia* infection in the control line was positively correlated with the adult emergence across *D. citri* generations (Table 3).

**Table 3.** Correlations among *Wolbachia* infection in *Diaphorina citri* and life history events from doxycycline-treated and control cultures of *D. citri* across nine consecutive generations.

| Doxycycline-treated line | Mean *Wolbachia* |  |  |  |  |
| --- | --- | --- | --- | --- | --- |
| (both sexes) | infection ($\Delta\Delta$ CT) | | | | |
| Life history event | correlation (*r*) | *R*<sup>2</sup> | *T* | *df* | *P* > *F* |
| Mean Female Proportion (%) | -0.67 | 0.44 | -2.68 | 9 | 0.025 |
| Control line |  |  |  |  |  |
| (both sexes) |  |  |  |  |  |
| Life history event | correlation (*r*) | *R*<sup>2</sup> | *T* | *df* | *P* > *F* |
| Mean Adult Emergence (%) | 0.83 | 0.70 | 4.54 | 9 | 0.001 |

### Sex ratio

Antibiotic treatment was a significant factor affecting the male : female sex ratio in the two *D. citri* treatment lines (Table 4). There was a significant female bias observed in the doxycycline-treated line as compared with the control during the F3 generation, and then during generations F5-F9 (Fig. 4). During the generations when a statistically significantly female bias was quantified, *Wolbachia* abundance was 54-89% lower in the doxycycline-treated line than in the control line. Overall, more female adults were produced by *D. citri* following doxycycline treatment than in the control. Additionally, mean *Wolbachia* abundance in the doxycycline-treated line was highly negatively correlated with the mean proportion of females produced during each generation of *D. citri* measured (Table 3).

**Figure 4.**
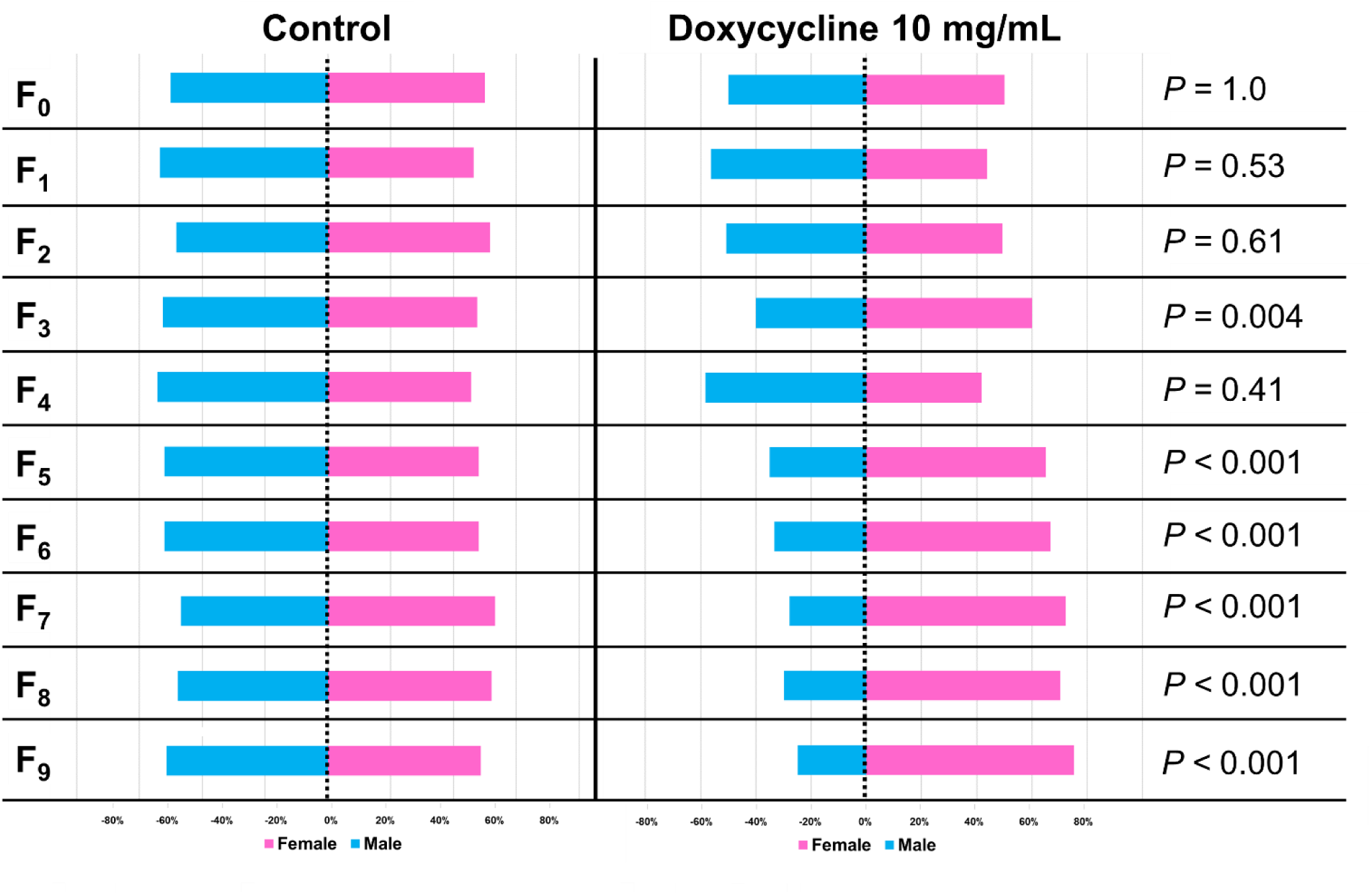
Effect of doxycycline treatment on *Diaphorina citri* sex ratios per generation. Chi-square, P < 0.05.

**Table 4.**
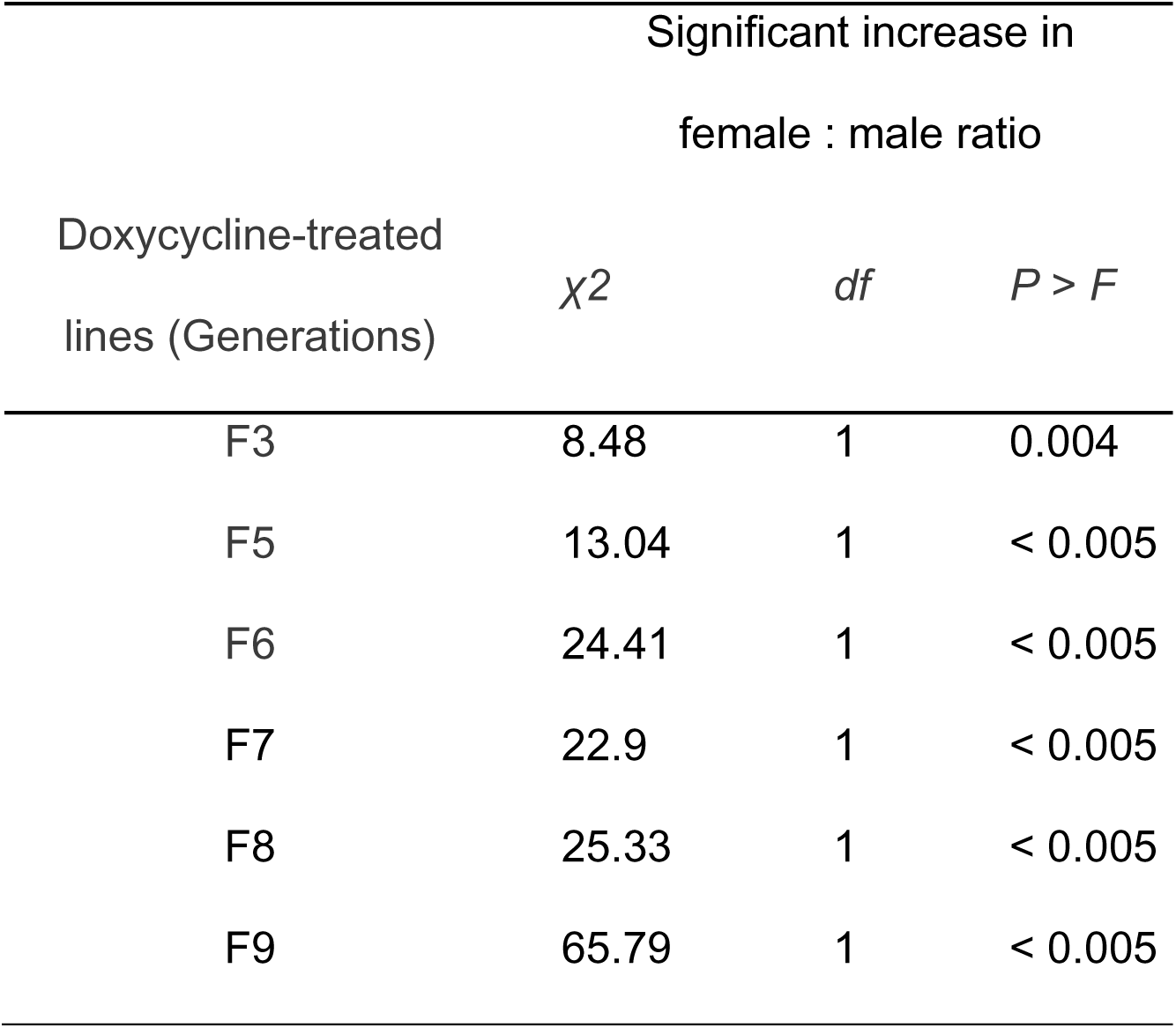
Statistical comparisons indicating a significant effect of doxycycline treatment on skewing the sex ratio of *Diaphorina citri* offspring towards female adults across nine consecutive generations.

### Longevity of survival

The effect of doxycycline treatment on longevity of *D. citri* adults was quantified during the F8 and F9 generations of the experiment. The survival probability of *D. citri* adults was significantly lower in the doxycycline-treated line than the control for F8 males (Log-rank test, χ2(1) = 101.46, P < 0.001) and females (Log-rank test, χ2(1) = 39.27, P < 0.001); as well as F9 males (Log-rank test, χ2(1) = 125.28, P < 0.001) and females (Log-rank test, χ2(1) = 18.01, P < 0.001) (Fig. 5A-D). Overall, survival of *D. citri* with reduced *Wolbachia* abundance was lower than that of control *D. citri*.

**Figure 5.**
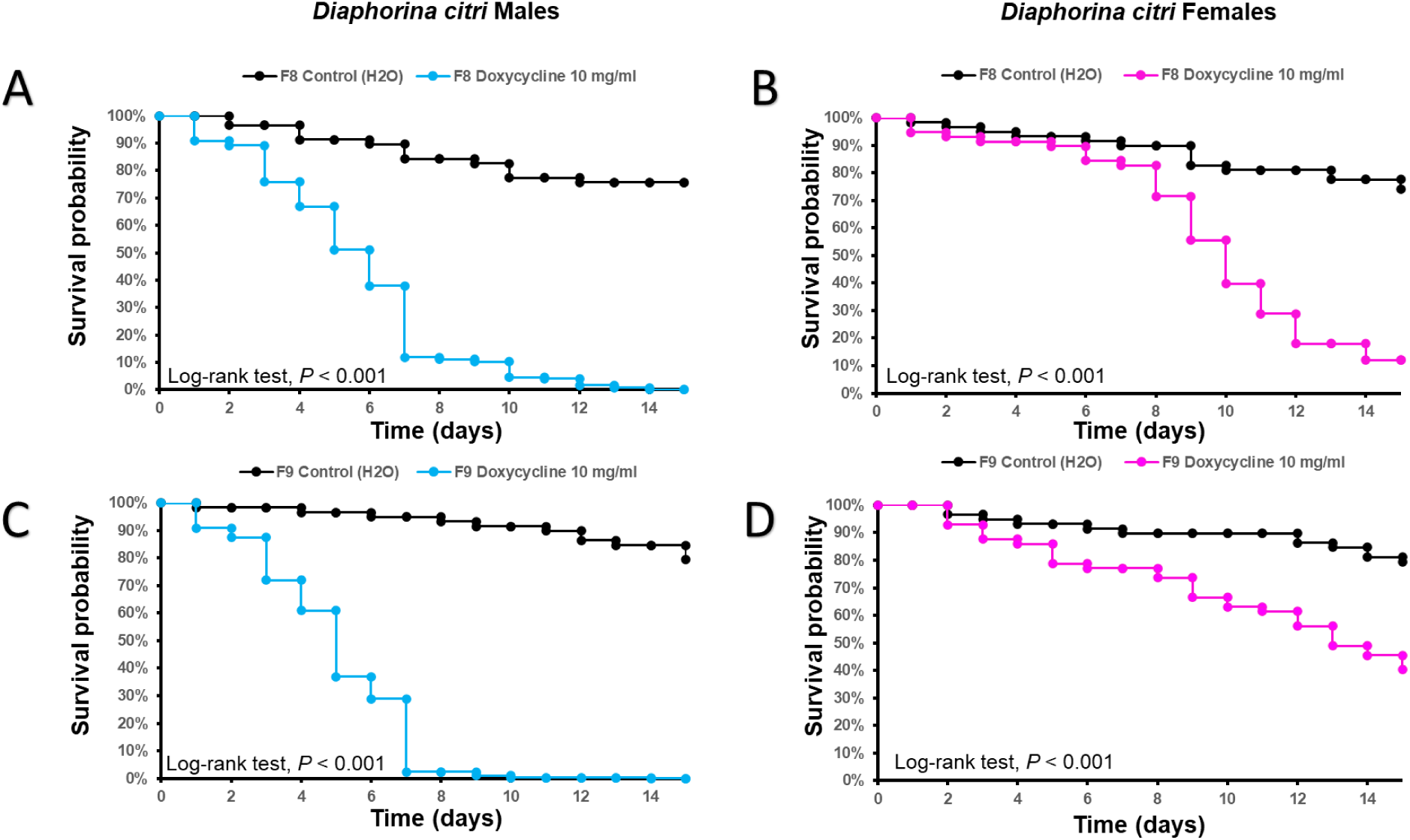
Effect of doxycycline treatment on survival of *Diaphorina citri* (A-C) males and (B-D) females during the F8 and F9 generations as displayed by Kaplan-Meier longevity curves. (A-D) Log-rank test, P < 0.05.

### *C*Las acquisition in *D. citri*

*D. citri* treated with doxycycline acquired significantly more *C*Las as both adults (AOV, F = 12.28, df = 2,1, P < 0.005) and nymphs (AOV, F = 5.63, df = 2,1, P = 0.004) than corresponding control psyllids during both the F8 and F9 generations (Fig. 6A-B). The *C*Las infection rate was significantly (Table 5) higher among adult *D. citri* from the doxycycline-treated line (77-80% *C*Las-positive) than in the control line (20-40% *C*Las-positive) (Fig. 6C). There were no differences (Table 5) in *C*Las infection rates between treatments in *D. citri* nymphs during the F8 (73-85% *C*Las-positive) and F9 (87% *C*Las-positive) generations (Fig. 6D). In addition, the abundance of *C*Las acquired by psyllids was negatively correlated with the abundance of *Wolbachia* infection in *D. citri* in the doxycycline-treated line (Table 6).

**Figure 6.**
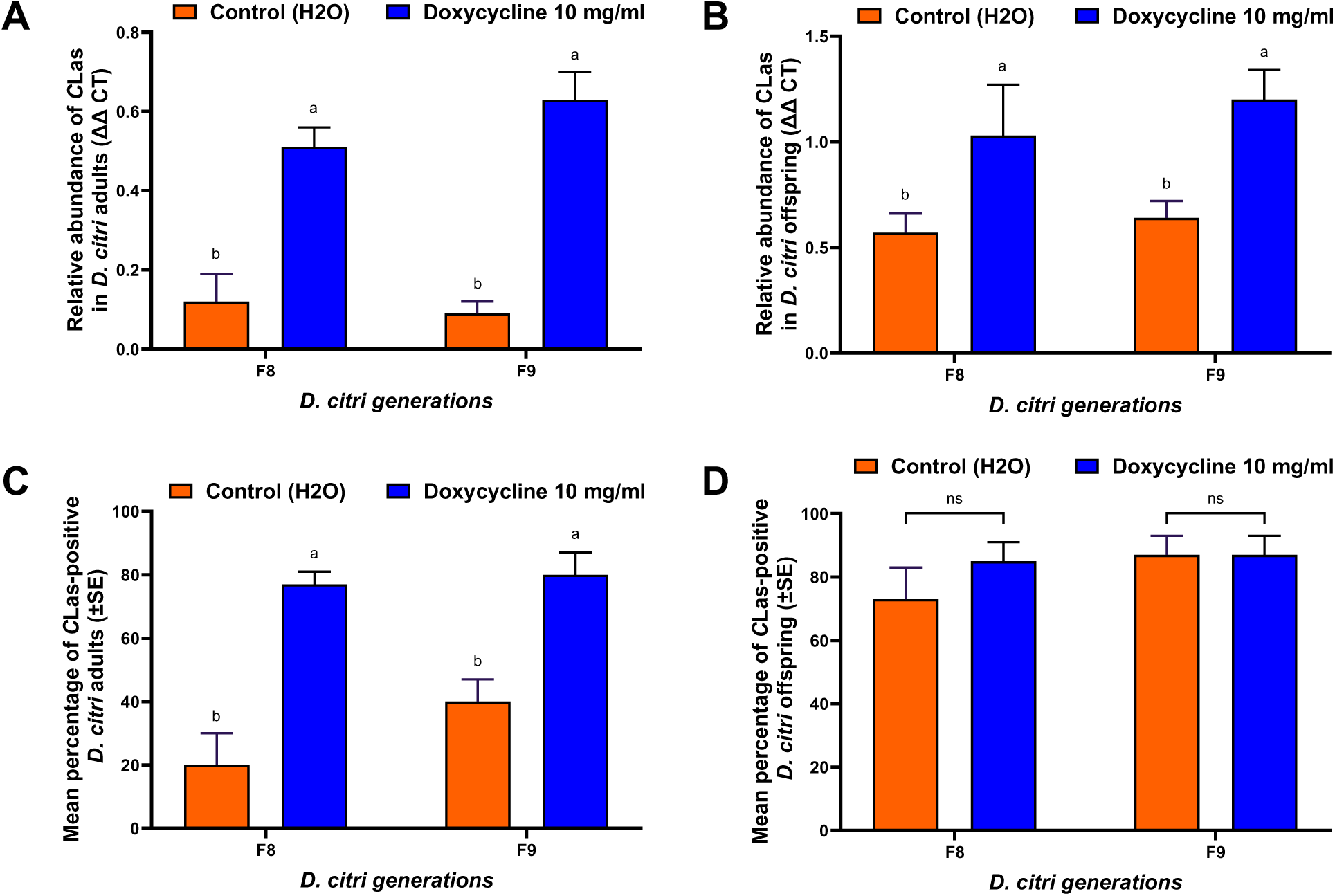
Effect of doxycycline treatment on acquisition of *C*Las by *Diaphorina citri* (A) adults and (B) offspring (nymphs), and infection rate of *D. citri* (C) adults and (D) offspring (nymphs) following acquisition assay on *C*Las-infected trees. Treatments indicated by “ns” are not significantly different. (A, B) ANOVA; P < 0.05; * < 0.1, Tukey’s HSD. (C, D) Chi-Square, P < 0.05.

**Table 5.** Statistical comparisons of the mean *C*Las infection rate in *Diaphorina citri* as affected by doxycycline treatment and generation of rearing.

| CLas infection rate in *D. citri* |  |  |  |  |  |
| --- | --- | --- | --- | --- | --- |
| Generation | Doxycycline-treated line | Control line |  |  |  |
| **Adults (both sexes)** | Mean IR (%) | Mean IR (%) | $\chi^2$ | *df* | *P > F* |
| F8 | 77 ± 4 | 20 ± 10 | 37.05 | 1 | < 0.005 |
| F9 | 80 ± 7 | 40 ± 7 | 15.19 | 1 | < 0.005 |
| **Offspring (both sexes)** | Mean IR (%) | Mean IR (%) | $\chi^2$ | *df* | *P > F* |
| F8 | 85 ± 6 | 73 ± 10 | 0.06 | 1 | 0.81 |
| F9 | 87 ± 6 | 87 ± 6 | 0.25 | 1 | 0.62 |

**Table 6.**
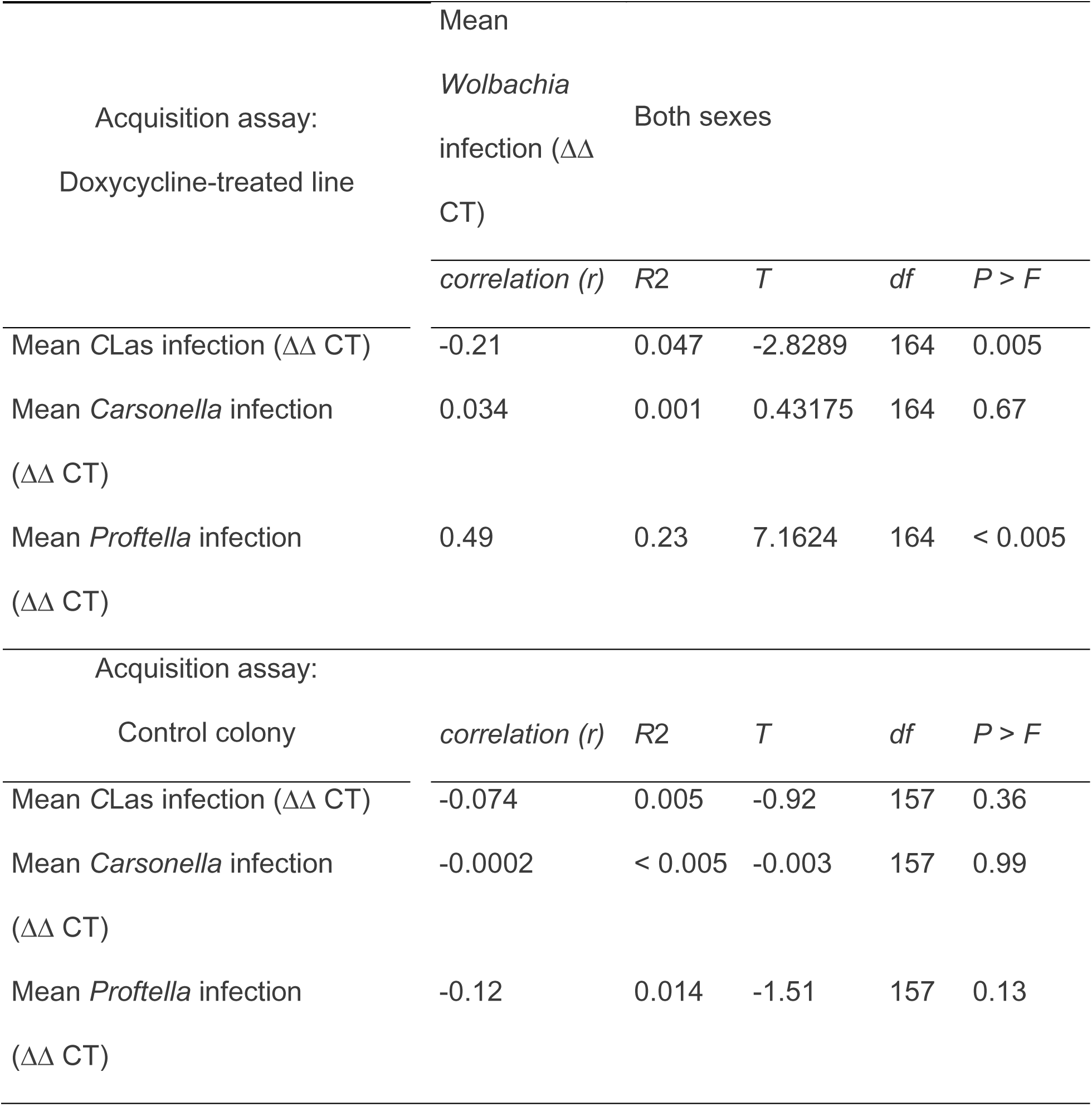
Correlations between *Wolbachia* infection and endosymbiont abundance after adults were exposed to *C*Las-positive plants for pathogen acquisition. Only adults from the F8 and F9 generations were used for this analysis.

There was no statistical difference in abundance of Carsonella between doxycycline-treated and control *D. citri* following acquisition of *C*Las by adults (AOV, F = 0.87, df = 2,1, P = 0.35). However, the abundance of Carsonella was lower in doxycycline-treated adults than in control adults during the F9 generation (Fig. 7A). There were no significant differences in Carsonella abundance in *D. citri* nymphs between treatments and among the generations tested (AOV, F = 0.04, df = 2,1, P = 0.96). Although the mean abundance of *Wolbachia* in doxycycline-treated *D. citri* appeared positively correlated with mean abundance of Carsonella, this correlation was not statistically significant (Table 6).

**Figure 7.**
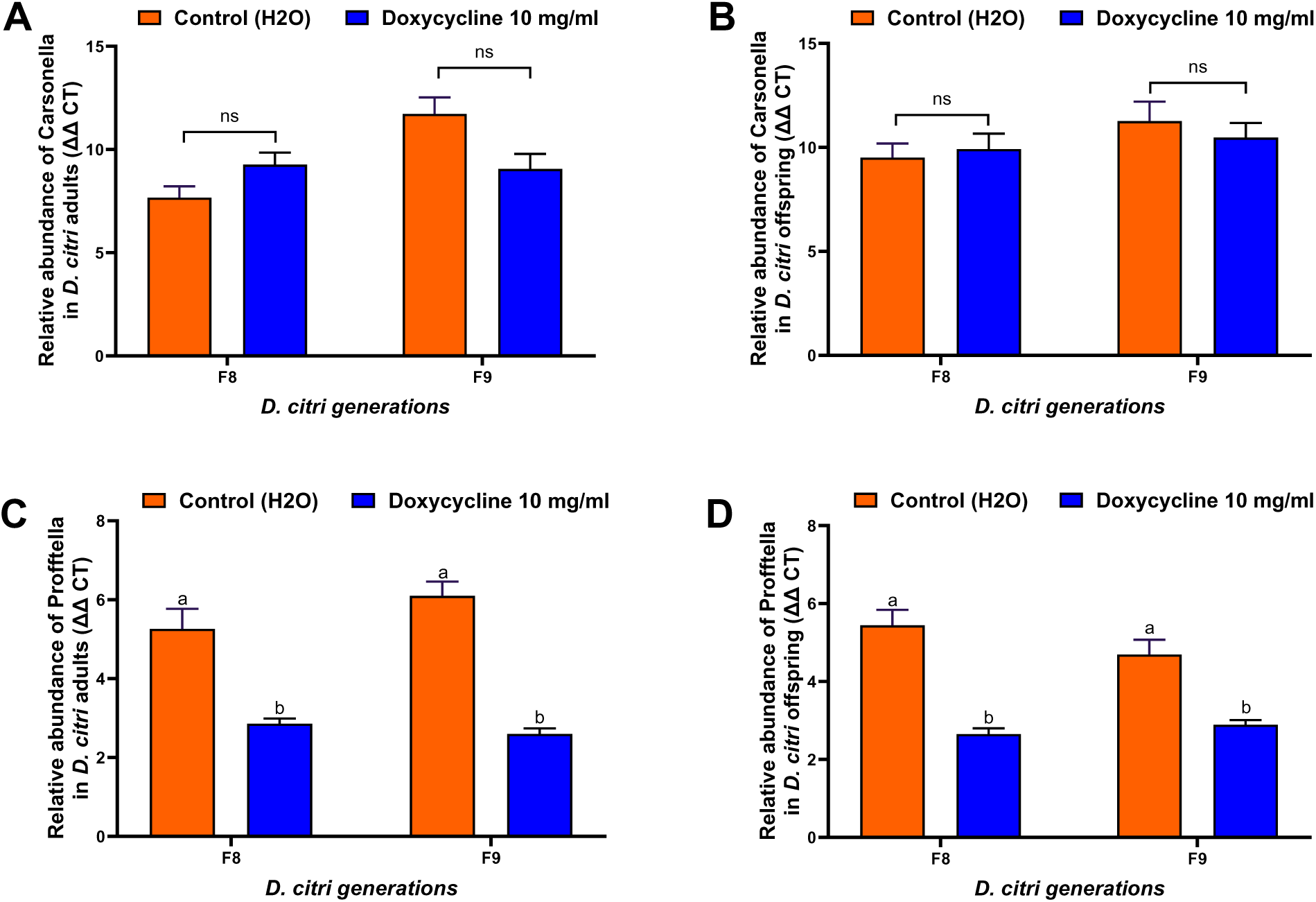
Effect of doxycycline treatment on Carsonella (A, B) and Proftella (C, D) abundance in *Diaphorina citri* adults and offspring. Treatments indicated by “ns” are not significantly different. (A-D) ANOVA; P < 0.05, Tukey’s HSD).

There were significant differences in abundance of Profftella between treatments and among generations of *D. citri* adults (AOV, F = 14.33, df = 2,1, P < 0.005) and offspring (AOV, F = 6.12, df = 2,1, P < 0.005). Abundance of Profftella was significantly lower in doxycycline-treated adults and nymphs than in the control line during both the F8 and F9 generations (Fig. 7C-D). In addition, the mean abundance of *Wolbachia* in doxycycline-treated *D. citri* was positively correlated with the mean abundance of Profftella (Table 6). In contrast, the mean abundance of *Wolbachia* in *D. citri* in the control line appeared negatively correlated with the mean abundance of Profftella, however, this correlation was not statistically significant (Table 6).

## Discussion

The purpose of this investigation was to eliminate or reduce the abundance of bacterial endosymbionts within a hemipteran phytopathogen vector, *D. citri*, with antibiotic treatment to gain insights into potential relationships between these organisms. *Diaphorina citri* is a piecing/sucking insect that vectors the pathogens causing citrus greening disease, which has severely limited commercial citriculture in several regions worldwide. Among the endosymbionts targeted, *Wolbachia* is particularly widespread among the Insecta and has profound effects on various aspects of general physiology, behavior, and ecology. To investigate the mechanisms by which endosymbionts affect insect biology often necessitates complete endosymbiont elimination from the host, followed by a comparison with infected isolines. In this investigation, *Wolbachia* load was reduced by 54% and 67% in both male and female *D. citri* after eight generations of antibiotic treatment. Furthermore, this reduction was observed to remain stable for two generations, although the extent of *Wolbachia* reduction exhibited variability across different generations. We believe this is the first successful demonstration of a substantial reduction in both *Wolbachia* and Profftella infections in *D. citri* using antibiotics. Our results also confirm that *Wolbachia* is associated with both fitness and female sex determination in *D. citri*, but also appears to compete with the *C*Las phytopathogen, which may be a target to exploit for disease management.

Antibiotic treatment in insects often yields diverse responses encompassing reductions in body mass, lifespan, relative growth rate, adult survival, embryo quality, altered feeding behaviors, and host selection (Prosser and Douglas, 1991; Douglas, 1992, 1995; Wilkinson and Douglas, 1995; Adams and Douglas, 1997). This study focused on fertility, measured by egg hatch rates, in doxycycline-treated *D. citri*. Surprisingly, the doxycycline-treated line exhibited higher fertility compared to controls; however, the control line produced significantly more offspring than the doxycycline-treated group. This contrasts with responses observed in other insect species, such as *Liposcelis tricolor*, where antibiotic treatment eliminated *Wolbachia* but did not cause a concomitant decrease in egg production or increase in embryonic mortality (Dong et al., 2006). In a separate investigation, high doses of tetracycline reduced wasp offspring production for long periods after treatment (Stouthamer and Mak, 2002; Lee et al., 2016).

Emergence of adult *D. citri* remained unaffected by antibiotic treatment in the current investigation. Interestingly, *Wolbachia* abundance was positively correlated with adult emergence in the control line, suggesting that this endosymbiont may facilitate development. Collectively, these results indicate mutualistic benefits of *Wolbachia* on reproductive fitness of *D. citri* more characteristic of an endosymbiont than that of a parasite. This observation mirrors the characteristics of many hemipteran host-endosymbiont associations. For instance, in the pea aphid, elimination of *Buchnera aphidicola* with antibiotic treatment reduced body mass of offspring, and decreased growth rate, survival, and embryo quality (Ohtaka and Ishikawa, 1991; Prosser and Douglas, 1991; Douglas, 1992, 1995; Wilkinson and Douglas, 1995; Adams and Douglas, 1997; Machado-Assefh et al., 2015). Similar effects have been documented across various insect genera, including Alydidae, Aphididae, Cimicidae, Culicidae, Drosophilidae, Dryinidae, Glossinidae, and Tephritidae (Dobson and Rattanadechakul, 2001; Pais et al., 2008; Brownlie and Johnson, 2009; Hosokawa et al., 2010; Lee et al., 2016; Espinosa et al., 2017; Wong et al., 2017).

Antibiotic treatment reduced abundance of *Wolbachia* in *D. citri* which was associated with a concomitant increase in the female-to-male sex ratio. The inverse relationship between abundance of *Wolbachia* and the proportion of female *D. citri* within the population indicates that the endosymbiont directly affected sex ratio but does not necessarily implicate cytoplasmic incompatibility as the causal mechanism. *Wolbachia* infections frequently lead to cytoplasmic incompatibility and male-killing in insects, resulting in populations with female-skewed sex ratios (Fialho and Stevens, 2000; Perlmutter et al., 2020). In the case of *Oryzaephilus surinamensis*, *Wolbachia* induces cytoplasmic incompatibility, ultimately serving to increase fitness of the bacteria (Kiefer et al., 2022). Egg hatch is reduced by 50% when *Wolbachia*-free *O. surinamensis* females, created via tetracycline treatment, are mated with *Wolbachia*-infected males; however, there are no indications that *Wolbachia* kills males in this species (Kiefer et al., 2022). In contrast, in the black flour beetle, *Tribolium madens*, *Wolbachia*-infected females consistently produced highly female-biased progeny, but the phenomenon is likely attributable to embryonic male-killing (Fialho and Stevens, 2000). Furthermore, *Wolbachia* infection is associated with reduced egg hatch rates in *T. madens*. Remarkably, tetracycline treatment effectively eliminated *Wolbachia* infection in *T. madens* and restored sex ratios to equilibrium levels, underscoring the dominant effect exerted by this endosymbiont on reproductive phenotype (Fialho and Stevens, 2000). It is important to note that our antibiotic treatment did not completely eliminate *Wolbachia* in *D. citri.* Although the level of reduction we achieved here greatly increased the proportion of females in the antibiotic-treated lines, this effect may be even greater if the endosymbiont were eliminated completely. If it is possible to do so in *D. citri* without an extinction even, the full extent of completely eliminating this endosymbiont from psyllids on resultant population sex ratio remains to be determined in this species.

There was considerable reduction in survival of both male and female *D. citri* subjected to doxycycline treatment as compared to their counterparts in the control line. By the F8 generation of the long-term isoline experiment, 90% of doxycycline-treated *D. citri* males and females were dead by the 7^th^ and 14^th^ day of observation, respectively. The results were similar in the F9 generation, where 95% of the doxycycline-treated males and 50% of treated females were dead by 7 and 14 days, respectively. In contrast, survival observed in the control line was above 80% at these same intervals (Fig. 5A-D). Therefore, abundance *Wolbachia* in *D. citri* appears to be strongly associated with lifespan and fitness. Congruent with our results, *Eretmocerus formosa* exhibit decreased lifespan following reduction of *Wolbachia* abundance as compared with control colonies (Wang et al., 2017).

Abundance of both *Wolbachia* and Profftella were substantially reduced in *D. citri* treated with doxycycline; however, the treatment did not affect abundance of the Carsonella endosymbiont. Furthermore, there was a positive correlation between abundance of *Wolbachia* with that of both Carsonella and Profftella in the untreated control line, suggesting potential interdependent relationships between these endosymbionts within *D. citri.* These results are consistent with both sex- and location-based correlations in abundance of these endosymbionts described previously in *D. citri* (Chu et al., 2016).

Our results indicate that doxycycline treatment had a more selective impact on abundance of *Wolbachia* than that of the other two endosymbionts quantified. Also, although doxycycline reduced *Wolbachia* populations in both sexes, there was a more pronounced reduction in females than in males. In addition, decreasing abundance of *Wolbachia* in *D. citri* increased acquisition of the *Candidatus* Liberibacter asiaticus (*C*Las) phytopathogen compared to that observed in controls where *Wolbachia* was not manipulated. These results indicate an inverse relationship between abundance of *Wolbachia* within *D. citri* and its competence as a vector of *C*Las and have potential implications for management of citrus greening disease. First, treatment of infected trees with broad-spectrum antibiotics, such as doxycycline, may inadvertently increase vector competence among populations of wild-type *D. citri* by reducing their natural *Wolbachia* populations. Alternatively, it might be possible to manipulate abundance of *Wolbachia* within natural populations of *D. citri* to reduce their vector competence. However, this will require a greater understanding of how *Wolbachia* may interfere with or at least diminish *C*Las acquisition. For example, it is possible that *Wolbachia* titers are only indirectly associated with vector competence in *D. citri*, which may occur via a yet undiscovered intermediary mechanism or effector.

Development of a *Wolbachia*-free line of *D. citri* holds promise to further unravel the dynamics between this endosymbiont and its host. However, the pursuit of an entirely *Wolbachia*-free population of *D. citri* within the confines of a laboratory setting is impeded by the substantial negative fitness consequences associated with elimination of this endosymbiont. Various strategies have been employed to eradicate *Wolbachia* from insects, including the use of antibiotic-treated hosts (Dobson and Rattanadechakul, 2001; Pais et al., 2008; Brownlie et al., 2009; Hosokawa et al., 2010; Lee et al., 2016; Espinosa et al., 2017; Wang et al., 2017), and heat treatment (Van Opijnen and Breeuwer, 1999). However, the effectiveness of symbiont removal with such methods varies and complete elimination of endosymbiont populations is not universally achievable (Smith and Rajan, 2000; Ruan et al., 2006; Wang et al., 2017; Gunderson et al., 2020). In this investigation, we were able to reduce populations of *Wolbachia* in *D. citri* considerably, but endosymbiont populations were not completely eliminated. A 60% reduction in abundance of *Wolbachia* achieved here had profound negative consequences on survival, development, and vector competence in antibiotic-treated *D. citri*, suggesting that endosymbionts may serve as useful targets in this vector to yield beneficial tools for management of citrus greening disease. Further attempts to develop an isoline of *D. citri* entirely devoid of *Wolbachia* may further benefit this pursuit.

## Acknowledgements

The authors are grateful to Christine Click, Rosa Johnson, and Alyssa Brown for their technical assistance. This work was funded in part by USDA NIFA agreement number 2021-70029-36053 to K.P.-S.

